# EstroGene database reveals diverse temporal, context-dependent and directional estrogen receptor regulomes in breast cancer

**DOI:** 10.1101/2023.01.30.526388

**Authors:** Zheqi Li, Tianqin Li, Megan E. Yates, Yang Wu, Amanda Ferber, Lyuqin Chen, Daniel D. Brown, Jason S. Carroll, Matthew J. Sikora, George C. Tseng, Steffi Oesterreich, Adrian V. Lee

## Abstract

As one of the most successful cancer therapeutic targets, estrogen receptor-α (ER/ESR1) has been extensively studied in decade-long. Sequencing technological advances have enabled genome-wide analysis of ER action. However, reproducibility is limited by different experimental design. Here, we established the EstroGene database through centralizing 246 experiments from 136 transcriptomic, cistromic and epigenetic datasets focusing on estradiol-treated ER activation across 19 breast cancer cell lines. We generated a user-friendly browser (https://estrogene.org/) for data visualization and gene inquiry under user-defined experimental conditions and statistical thresholds. Notably, documentation-based meta-analysis revealed a considerable lack of experimental details. Comparison of independent RNA-seq or ER ChIP-seq data with the same design showed large variability and only strong effects could be consistently detected. We defined temporal estrogen response metasignatures and showed the association with specific transcriptional factors, chromatin accessibility and ER heterogeneity. Unexpectedly, harmonizing 146 transcriptomic analyses uncovered a subset of E2-bidirectionally regulated genes, which linked to immune surveillance in the clinical setting. Furthermore, we defined context dependent E2 response programs in MCF7 and T47D cell lines, the two most frequently used models in the field. Collectively, the EstroGene database provides an informative resource to the cancer research community and reveals a diverse mode of ER signaling.

## Introduction

More than two-thirds of breast cancers express estrogen receptor-α (*ER/ESR1*)(1) and therapeutic strategies blocking ER signaling is a long-standing and effective treatment strategy (2,3), though the landscape of breast cancer treatments is constantly evolving, such as the recently incorporated immune checkpoint inhibitors(4,5). Unfortunately, resistance to hormonal therapy remains a barrier and a large public health issue (6,7). Numerous mechanisms of resistance have been uncovered including genetic alterations in ER action, hotspot mutations(8,9), fusions(10) and *ESR1* copy number amplification(11). A thorough understanding of ER action in breast cancer is a key breast cancer research goal, which could pave a path towards novel ER-target therapies.

As a member of nuclear receptor family, ER is vital in sensing external hormonal cues and triggers various downstream phenotypic cascades in breast cancer cells. Classically, upon activation by ligands, ER forms dimers and binds to sites on DNA to enhance gene expression(12,13). In addition, recent studies have discovered several alternate ER effects such as modifying 3D chromatin loops to bring genes together for coordinated transcriptional regulation(14), modifying epigenetic factors such as FOXA1 to reshape chromatin landscapes(15) and controlling mRNA metabolism to sustain cell fitness towards external stressors(16). The ER signaling cascade is extraordinarily dynamic, heterogenous and context-dependent. For instance, a recent study showed that prolonged E2 administration induces transcriptional output and chromatin landscape fluctuations partially attributed to H2A ubiquitin ligase RING1B(17). Further, single-cell multi-omics delineated two E2 response programs associated with ER and FOXM1 respectively, which designates distinct chromatin accessibility states(18). The complex nature of the ER regulatory machinery provides a challenge to its dissection and understanding.

Advances in sequencing technologies have evolved at an unprecedented rate during the past decade and have largely facilitated genome-wide profiling of ER action in breast cancer(19). For example, RNA-sequencing together with earlier probe-based microarray platforms revealed genome-wide E2-induced transcriptomic changes (20). Likewise, ER ChIP-sequencing identified differential ER binding regions (21). The rapid development of single-cell omics will provide greater granularity and allow assessment of the heterogeneity of E2 response (18). Nevertheless, the benefits of the rapidly growing data sets of ER action, many of which are publicly available, is limited in part due to the paucity of an end-to-end data harmonization for uniform data curation, processing, and analysis.

Researchers have several data sets to choose from when examining if a gene of interest is regulated by E2, albeit repeated analysis of individual datasets consumes time and often reveals experimental variation and lack of reproducibility. Inter-data set variations are expected even under the same design due to the potential for different cell lines source, reagents, and sequencing platforms, and thus an E2-related multi-omic database which is comprehensive in its inclusion of available datasets is in great demand. We therefore developed the EstroGene knowledgebase to overcome these barriers. Unlike other databases such as the Cistrome DB(22) and GREIN(23), which primarily focus on providing access to data from a single omic platform and without regards to a specific type of experimental analysis, or other broadly-targeted nuclear receptor omic database such as the Transcriptomine (24) and Signaling Pathways Ominer (25), EstroGene focuses on a simple E2 stimulation experimental design in breast cancer cell lines and integrates multiple types of data covering transcriptome, genomic occupancy and chromatin interaction profiling. EstroGene provides a user-friendly browser allowing researchers a fast and comprehensive overview of ER regulation and concordance across hundreds of curated experiments. Once developed, we used the diversity of experimental conditions (i.e., E2 stimulation duration, doses, and models) to perform an in-depth analysis to elucidate the directionality of E2 response, temporal, trajectory, and contextual dependencies. We believe that EstroGene provides a useful analytic tool to help researchers rigorously and efficiently accelerate new discoveries on estrogen receptor biology in breast cancer and will ultimately facilitate the development of novel therapeutic concepts for treatment of endocrine resistance in breast cancer.

## Results

### Ingestion, annotation, and curation of sequencing data from estradiol stimulated breast cancer cells

To collect publicly available ER-related sequencing data sets, we initiated this project by mining data from the Gene Expression Omnibus using the key words “estrogen” or “E2” or “estradiol” plus “breast cancer” plus “the name a specific type of sequencing technology” (e.g., “RNA-seq” or “RNA-sequencing”). Our search strategy included seven widely used sequencing techniques including transcriptomic profiling (RNA-seq, microarray, GRO-seq), genomic occupancy profiling (ER ChIP-seq), chromatin accessibility profiling (ATAC-seq) and chromatin interaction profiling (ER ChIA-PET and Hi-C). We focused on estradiol (E2) stimulation in charcoal stripped and/or serum-free treated breast cancer cell lines, but also included limited ChIP-seq datasets on ER action in complete medium. Results were further manually filtered to ensure the corresponding study was suitable. To extend the database, we are further crowdsourcing datasets via social media and a google form (https://docs.google.com/spreadsheets/d/1PFMGB_-COSrUujMKl_M-Ogkmq4cAyuIEgoLwSLFuYTs/edit#gid=0) (Fig. 1A and Supplementary Table S1).

**Figure 1.**
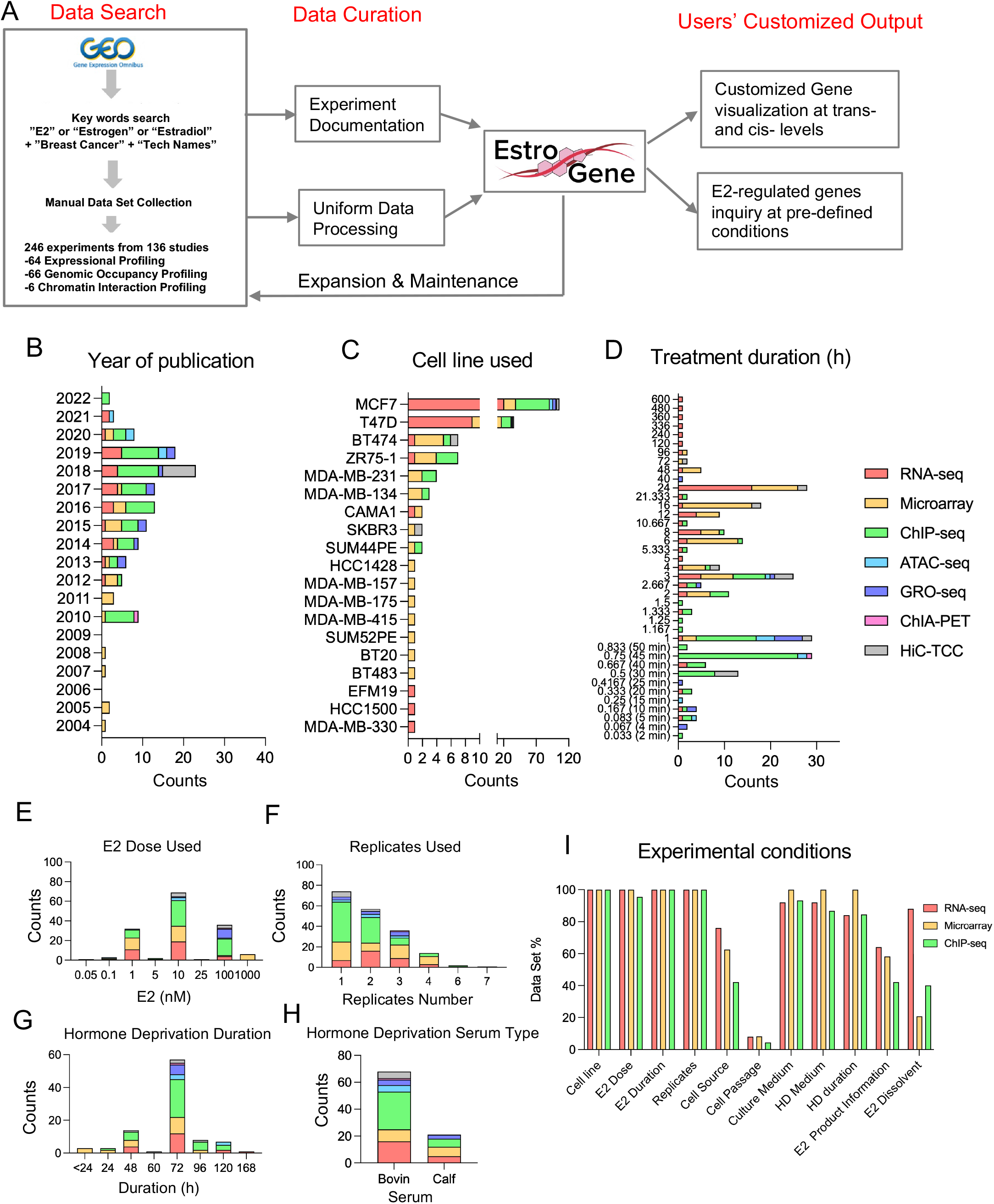
Ingestion, annotation, and curation of sequencing data from estradiol stimulated breast cancer cells. A. A flow chart depicting the process for establishment of the EstroGene database and specify the embedded functions of the browser. B to G. Stacked histogram showing the metadata separated by technologies across all the curated data sets related to year of data set publication (B), cell line used (C), E2 treatment duration (D), E2 dose selection (E), replicates used (F), hormone deprivation duration (G) and serum type (H). I. Bar graph showing the percentage of RNA-seq, microarray and ChIP-seq data sets with available detailed experimental terms.

We curated a total of 136 different datasets including 64 expression, 66 genomic occupancy and 6 chromatin interaction profiling studies published from 2004 to 2022 (Table 1 and Fig. 1B). Of note, a large portion of these experimental designs included multiple cell lines, E2 doses or time points, which resulted in 246 individual experimental conditions.

**Table 1.** Overall data set and data point included in the EstroGene database. Studies stand for independent data sets with individual GEO accession numbers. Experiments stand for a analytic comparison of E2 effects within one cell model or a single ER ChIP-seq profiling in full medium condition. Each study possibly contains several different experiments.

| Data Type | Technique | Studies | Experiments |
| --- | --- | --- | --- |
| Expressional Profiling | RNA-seq | 25 | 66 |
|  | Microarray | 29 | 80 |
|  | GRO-seq | 10 | 10 |
| Genomic Occupancy Profiling | ER ChIP-seq | 61 | 75 |
| Chromatin Accessibility Profiling | ATAC-seq | 5 | 6 |
| Chromatin Interaction Profiling | ER ChIA-PET | 1 | 1 |
|  | Hi-C/TCC | 5 | 8 |
| Total |  | 136 | 246 |

Chronologically, the number of sequencing datasets increased after 2010 (96.1% of all datasets) whereas microarray was the only technique used for RNA expression analysis before 2010. Transcriptomic (RNA-seq and microarray) and ER genomic occupancy profiling (ChIP-seq) were the most frequently applied methods (81.3%) (Fig. 1B), suggesting that the current understanding of ER action in breast cancer still mainly relies on the classic cistrome-to-transcriptome regulation. MCF7 and T47D cell lines accounted for nearly 80% of all experiments (Fig. 1C) (Fig. 1C). The duration of E2 exposure was mostly depended upon the investigation: transcriptomic profiling typically used a longer duration (69.5% above 6 hours) while cistrome profiling mainly captured a more rapid E2 response (71.1% below 1 hour) (Fig. 1D). All studies used saturated doses of E2, with 10 nM as the most frequently chosen dose followed by 1 nM and 100 nM (Fig. 1E). Approximately 30% of transcriptomic and 50% of ER cistromic profiling did not include biological replicates (Fig. 1F). In addition, hormone deprivation prior to E2 exposure varied with 61% of experiments performing hormone deprivation for 72 hours followed by 48 hours(Fig. 1G) and 76% studies using charcoal-stripped fetal bovin serum rather than calf serum (Fig. 1H).

We hand-abstracted methodological details from the original publications or GEO profiles to determine to what extent experimental conditions were reported. Data-set level documentations such as source publications, experimental conditions and sequencing parameters are presented in Supplementary Data Table S1. We focused on RNA-seq, microarray and ChIP-seq as these studies had the greatest number of datasets. Essential experimental terms such as cell line name, E2 doses and treatment durations were included in nearly all data sets, however, specific details were frequently missing. Sources of cell lines, which has been stated as a pivotal cause of inter-dataset inconsistency(26), was only included in 42% of the studies. Cell passage number, an important indication of cell state(27), was missing in 96% of these reports. And finally, sources of estradiol used and their corresponding diluent were missing in 60% of documentations (Fig. 1I).

### EstroGene: a multi-omic database of ER-regulated action

To enable researchers lacking sequencing and/or bioinformatics skills, we developed a web server named EstroGene (https://estrogene.org/) for data access and visualization. We first downloaded and used a single pipeline for processing and analysis of the majority of curated publicly available transcriptomic (23 microarray and 25 RNA-seq) and genomic occupancy data sets (32 ER ChIP-seq). The Estrogene web browser home page consists of a general introduction to this project and highlights various features with the corresponding hyperlinks. Two major browsing modules are embedded to fulfil different research purposes: a user-defined single gene-based data visualization function and a user-defined statistical cutoff-based gene list query function.

First, users can generate volcano plots and visualize the concordance of gene expression changes induced by E2 treatment from universal processing of 80 microarray and 66 RNA-seq individual comparisons across 17 and 8 breast cancer cell lines, respectively. In addition, users can limit the search by cell line, E2 dose and treatment duration. We present the percentage of comparisons showing up- or down-regulation of a certain gene (adjusted p value<0.05 for comparisons with replicates or |Log2FC|>0.5 for comparisons without replicates), to help users quantitatively evaluate the trend and consistency of regulation. For instance, the well-characterized estrogen-induced gene *GREB1* shows up-regulation in ~70% of microarray and RNA-seq analyses (Fig. 2A), while the estrogen-repressed gene *IL1R1* shows down-regulation in 47% (Fig. 2B). In contrast, the house-keeping gene *GAPDH* displayed a minimal degree of changes (with very few exceptions) across all the comparisons (Fig. 2C). The plots can be directly exported as a JPEG format file for further presentation. Of note, users can also click on each data point to access the original GEO submission for easy access and further analysis of the experimental details of certain data sets.

**Figure 2.**
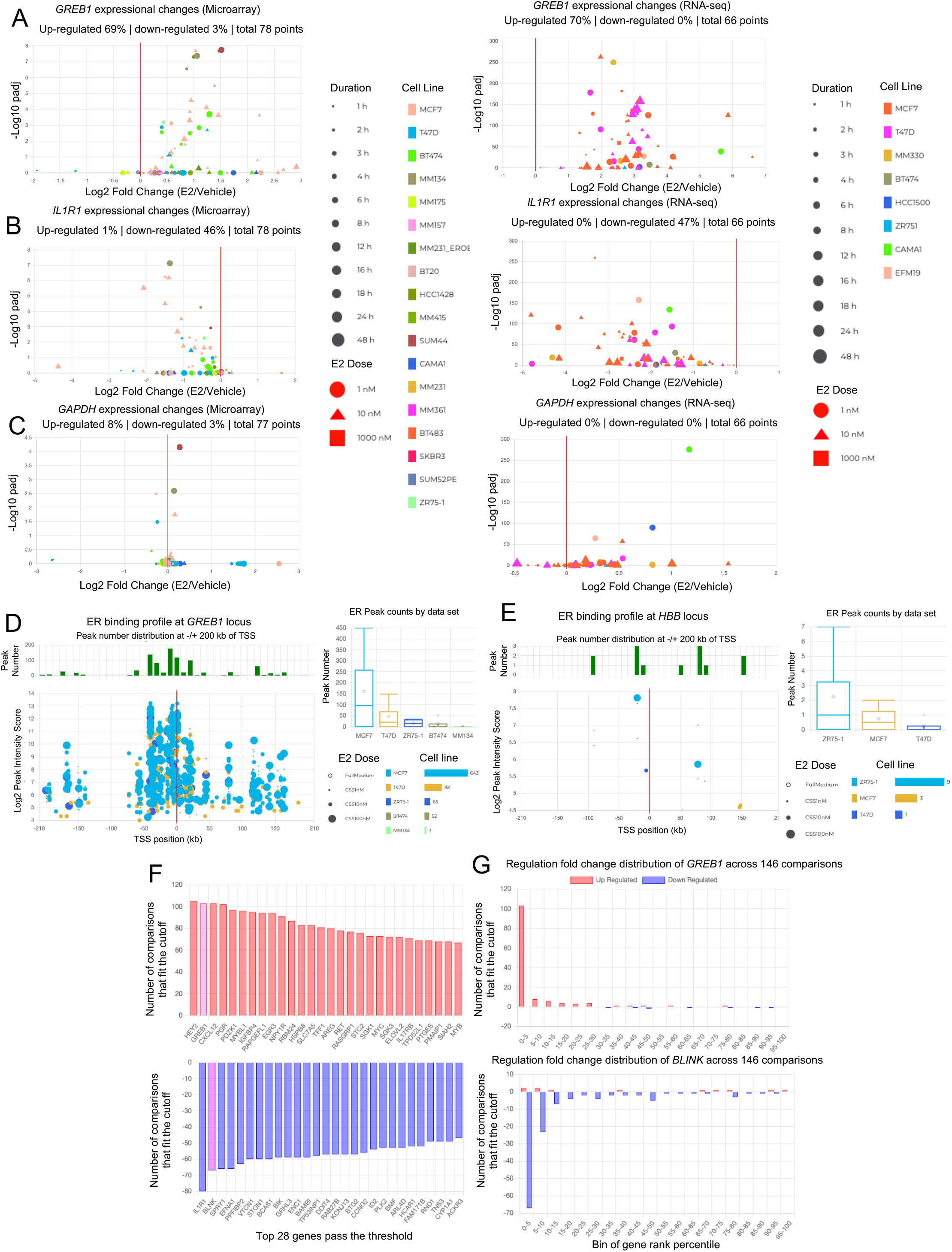
EstroGene: a multi-omic database of ER-regulated action. A to C. Screen shots from the EstroGene browser representing the transcriptomic consistency plots from RNA-seq (Left panel) and microarray (Right panel) towards a E2-induced GREB1 (A), a E2-repressed gene IL1R1 (B) and a non-E2 regulated gene GAPDH (C). D and E. Screen shots from the EstroGene browser showing the ER ChIP-seq consistency plots of the upstream and downstream 200kb of TSS towards a E2 regulated gene *GREB1* (D) and non-E2 regulated gene *HBB* (E) as positive and negative controls respectively. F and G. Screen shots from the EstroGene browser showing the statistical cutoff-based gene list query function. Examples are for the top 30 output with the thresholds of top 5 percentile of up (F) and down(G) regulated genes and consistent across at least 20% of comparisons (Left panel). One of gene from each section (GREB1 and BLNK) is further selected for cross-dataset tendency visualization (Right panel)

Besides transcriptomes, the analysis page also presents ER binding sites (as determined by ER ChIP-seq) with the corresponding intensities within −/+ 200 kb range of the transcriptional start site (TSS) of each gene. The plot shows a combination of 32 uniformly processed ER ChIP-seq data sets in either full medium or E2-treated condition and can be filtered based on users’ defined condition. This feature is exemplified by the ER proximal binding landscape of *GREB1* gene which shows 954 peaks as a positive control (Fig. 2D), while only 13 binding sites were detected at the heterochromatin-enriched gene *HBB* serving as negative control (Fig. 2E).

In addition to the visualization module, we also provide a gene-list query function shown in the “Statistics” page that can be found on the home page. This module is based upon calculation of the percentile rank of each gene after merging 146 comparisons from RNA-seq and microarray. Users can define specific cut-offs for 1) trend and intensity as a percentage of up- or down-regulation among all genes and 2) inter-data set consistency as a percentage of all comparisons. The output shows all genes fitting the cut-offs from the highest to lowest consistency (Fig. 2F) and can be customized based upon the desired conditions and contexts. Importantly, users can plot the pattern of individual gene regulation across all the selected comparisons (Fig. 2G).

### Inter-dataset concordance of estrogen-induced transcriptomics

Given that most studies on ER rely on results gained from a single data set, we used EstroGene to address variation amongst different studies and assessed the concordance from experiments using the same conditions. For this analysis, we focused on RNA-seq and ChIP-seq data as they are the most abundant types of profiling.

We selected four independent RNA-seq experiments in MCF7 cells with each having aa very similar design of 24 hours of 10 nM E2 exposure and at least two biological replicates(28–31). Principle component analysis (PCA) revealed the greatest difference to be technical and associated with the individual dataset (Fig. 3A, left panel). This was to be expected as some minor technical variations in cell culture, hormone deprivation and sequencing platform were noted (Supplementary Table S2). This difference was mostly eliminated by batch effect correction, which showed the major variation to be alterations in the E2-induced transcriptome (Fig. 3A, Right panel). Of note, dataset R5 showed a greater difference in the E2-regulated transcriptome possibly due to the longer duration of hormone deprivation (168 hours) compared to the others (72 hours). This was also in line with numbers of differentially expressing genes (DEGs) computed under four different fold change thresholds (Fig. 3B). The number of replicates used in the analysis did not correlate with the number of DEGs.

**Figure 3.**
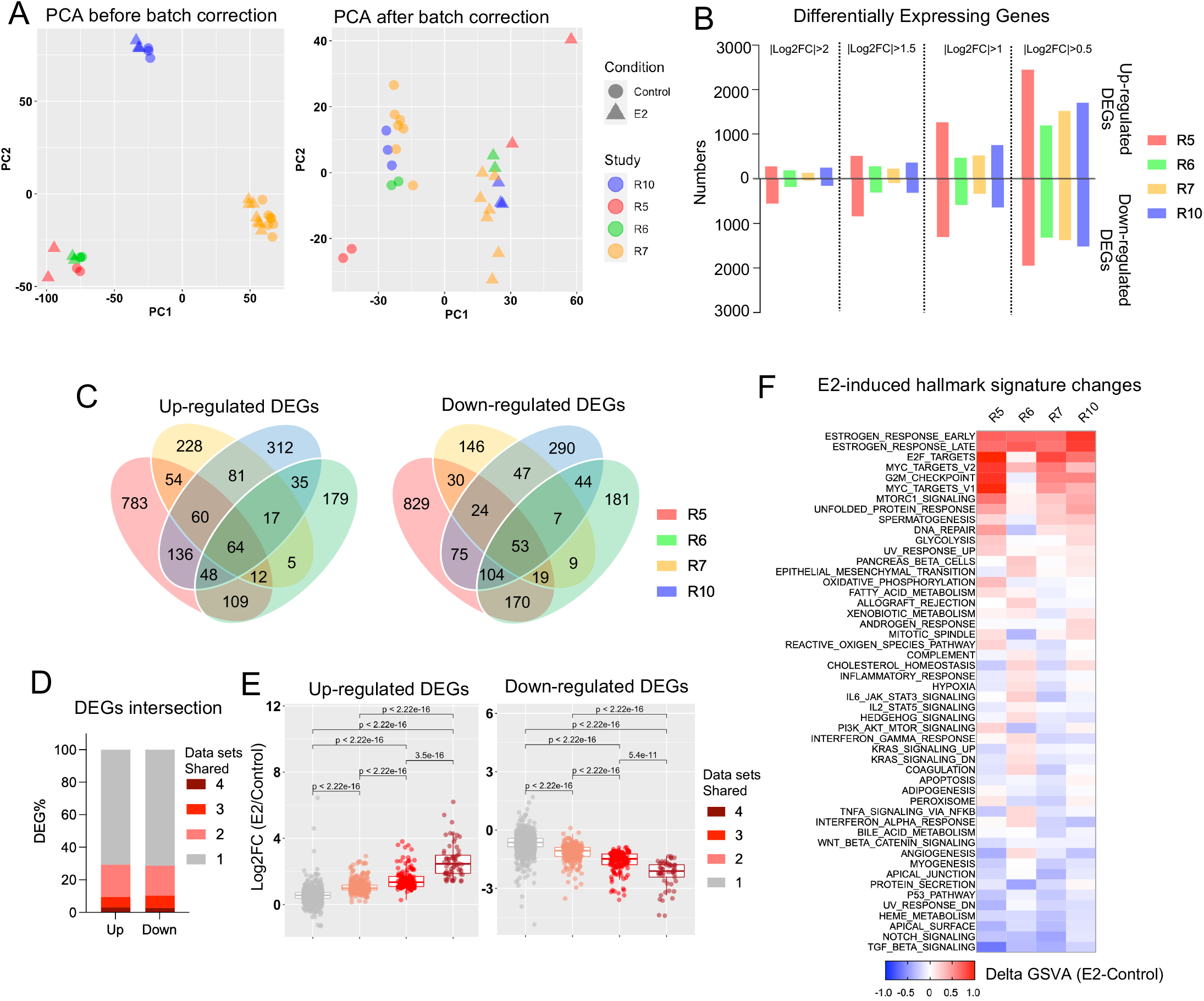
Inter-dataset concordance of estrogen-induced transcriptomics. A. Principal component analysis depicting the cross-sample variation from four independent RNA-seq experiments before (Left panel) and after (Right panel) batch effect correction. B. Bar chart showing the number of up- and down-regulated differentially expressing genes from the four RNA-seq comparisons with under four different fold change cutoffs. C. Venn diagrams depicting the overlap of E2-induced up (Left panel) and down (Right panel) regulated differentially expressing genes from the RNA-seq analysis using a cutoff of |log2FC|>1 and padj<0.05. D. Stacked plot showing the percentage of DEGs shared in one to four data sets. E. Box plot showing average log2 fold changes from four RNA-seq experiments of each up (Left panel) and down (Right panel) regulated genes from the four different consistency classes. Mann Whitney U test was used. F. Heatmap summarizing the E2-caused enrichment score differences of 50 Hallmark gene sets across four RNA-seq experiments.

The intersection of up- and down-regulated DEGs from four data sets (|log2FC|>1, padj<0.05) showed that greater than 70% of DEGs were unique to one experiment, whereas only ~3% genes were regulated in all four data sets (Fig. 3C and 3D, Supplementary Table S3). We classified the DEGs into four classes based upon the number of data sets they shared and compared the fold change regulation (Supplementary Table S3). The consistency of both up- and down-regulation of genes was strongly correlated with the degree of E2-regulation (Fig. 3E), with moderately or weakly E2-regulated genes being uniquely regulated in individual experiments. In addition, comparison of the enrichment of E2-induced genes in hallmark signatures showed considerable variation for most of the pathways except the well-characterized estrogen response signatures and proliferation-related signatures (Fig. 3F). For instance, the cholesterol homeostasis signature enrichment showed either a modest increase or decrease after E2 stimulation in two of the comparisons. Of note, data set R6 showed a more distinct pathway alteration pattern compared to the other three, which might be partially owning to the use of fetal calf serum rather than fetal bovine serum in maintenance and hormone deprivation. In summary, we demonstrate that strong transcriptomic changes are reproduced across multiple data sets, but moderate-to-weak E2-induced changes are often inconsistent between different data sets and thus caution may be needed when interpreting results from a single experiment or data set.

### Inter-dataset concordance of estrogen-induced ER genomic binding

We investigated the similarity of E2-regulated ER chromatin binding profiles among four different ER ChIP-seq data sets generated from MCF7 cells treated with 10 nM E2 for 45 minutes after 48 or 72 hours of hormone deprivation(32–35). Importantly, the same ER antibody (Santa Cruz sc543) was utilized in all experiments to pull down ER (Supplementary Table S4). Like transcriptomic data processing, all raw sequencing files were aligned, and peaks were called using an identical pipeline. We also examined the quality parameters (e.g., percentage of reads within peaks and percentage of reads within Blacklist regions) of derived peaks to examine technical variation (Supplementary Table S5). The number of peaks in the vehicle groups showed a large variation ranging from 332 (C42) to 23,189 (C42) (Supplementary Table S5). This difference may in part be due to a shorter hormone deprivation duration (48 hours) in C42 which shows the largest number of baseline ER binding. As expected, E2 stimulation resulted in a 1.6-to-27.5-fold gain of ER binding (Supplementary Table S5).

PCA revealed divergence among E2-treated samples whereas control samples mostly clustered together (Fig. 4A). Since three out of four data sets did not include replicates, we used an occupancy-based strategy to identify differential ER peaks (i.e., directly comparing control and E2 peaks). We observed 25,109 (C22), 20,260 (C33), 15,153 (C42) and 1,690 (C34) gained ER peaks with E2 treatment, whereas less than 70 peaks were lost (Fig. 4B). Notably, despite the discrepancy in the number of gained ER peaks, the peaks showed a similar distribution of genomic features such as promoter and distal intergenic regions (Supplementary Fig. S1A), suggesting that there was not a genomic location bias associated with the biological variation. Like the transcriptomic analysis, intersecting gained ER peaks across the four datasets showed limited consistency: approximately 60% of peaks were only exclusively present in a single data set while merely 1.3% peaks (N=519) were shared in all four data sets while, again highlighting the high degree of inter-dataset discordance (Fig. 4C and 4D).

**Figure 4.**
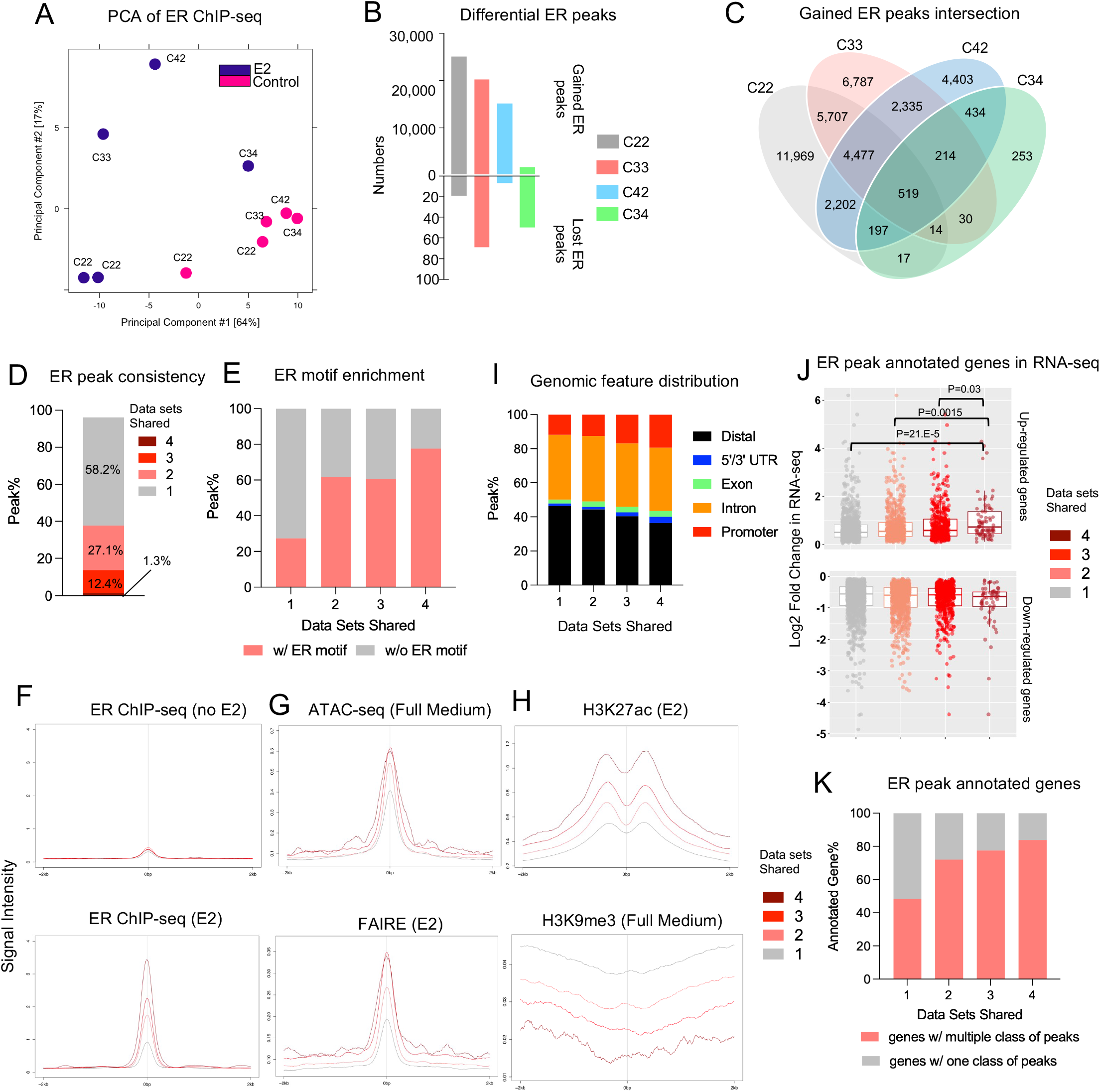
Inter-dataset concordance of estrogen-induced ER genomic binding. A. Principal component analysis depicting the ER genomic binding variations across four different experiments. B. Bar plot showing the number of gained and lost ER peaks from four ChIP-seq experiments. C. Venn diagram showing the intersection of E2-induced gained ER peaks across four ChIP-seq experiments. D. Stacked bar plot depicting the percentage distribution of gained ER peaks consistent in one to four experiments. E. Stacked plot representing the percentage of peaks containing ER motif across four peak sets. F to H. Intensity plot showing the binding signals from ER ChIP-seq in the presence or absence of estrogen (F), ATAC-seq and FAIRE (G) and H3K27ac/H3K9me3 ChIP-seq at the four gained ER peak sets with different cross-data set consistencies. ER ChIP-seq and epigenetic profiling data sets were downloaded from GSE78284, GSE25710, GSE102441, GSE78913 and GSE96517. I. Stacked plots showing the genomic feature distributions of the four gained ER peak sets with different cross-data set consistencies. J. Box plots depicting the average log2 fold changes from four RNA-seq experiments in Fig. 3 towards all the up (Top panel) and down (Bottom panel) regulated genes annotated from −/+ 50 kb of the four gained ER peak sets. Mann Whitney U test was used. K. Stacked plot representing the percentage of annotated genes in J with one or multiple consistency class of gained ER peaks.

We annotated all E2-induced ER peaks based upon the number of data sets they were found in where “peak set 4” represents consistently gained ER peaks in all four datasets, and “peak set 1” stands for those only found in a single dataset. A motif scan demonstrated that highly consistent gained ER peaks (peak set 4) were more likely to be enriched in ER motifs (77.6% in peak set 4) compared to those low-consistent gained ER peaks (27.3% in peak set 1) (Fig. 4E). Analysis of peak intensity also showed that consistent peaks in general exhibited stronger binding intensity upon E2 stimulation, whereas no differences was noted at baseline level (Fig. 4F). Integrating with publicly available epigenetic profiling from MCF7 cells in full medium or E2-stimulated conditions, peak set 4 was enriched in more active higher accessibility chromatin (Fig. 4G) as shown by increased H3K27ac and decreased H3K9me3 marks (Fig 4H). Finally consistent peaks were associated with increasing occupancy at promoter regions and decreasing occupancy at distal intergenic regions (Fig. 4I).

Annotation of genes associated with the ER ChIP-seq data sets revealed that the number of genes reached a plateau if we chose 50kb or a longer peak flank distance for the annotation (Supplementary Fig. S1B) (i.e., annotating genes from upstream and downstream of a certain distance from each peak). We thus used 50kb as the annotation distance for downstream comparison. When integrated with expression fold changes of the four RNA-seq data sets analyzed above, the genes associated with the gained ER peaks shared in all four data sets displayed significantly greater levels of mRNA induction compared to genes induced in the three other peak sets (Fig. 4J). In addition, highly consistent ER peaks are more likely to harbor ER binding sites also found in other other peak categories (Fig. 4K). Taken together, our analysis showed that highly consistent E2-induced ER binding sites among different data sets represent a subset of peaks with stronger E2-inducibility due to more direct ER binding potential and more accessible chromatin, and links to more pronounced transcriptional alterations because of a higher degree of promoter distribution and cooperativity with more other ER binding events.

Our inter-data set comparison of transcriptomic and cistromic regulation uncovered a high level of dissimilarity across different experiments, and only biological effects with the greatest effect sizes were conserved. To interrogate inter-data set consistency, we identified gene expression changes across 66 RNA-seq and 80 microarray experiments based upon the E2-induced fold change in gene expression. We derived the percentile of gene expression change for each individual gene normalized to all genes expression changes within each experiment, and filtered out genes that were detected in less than 80% of experiments (Supplementary Fig. S2A). First, each gene’s fold change percentile showed a positive correlation between the two platforms (Supplementary Fig. S2B), indicating that the vast majority of genes do not bear platform-based bias. We next derived consistency-to-inducibility maps consisting of 12,429 genes across 146 comparisons (Supplementary Fig. S2C). In line with our inter-data sets analysis above, highly inducible, or repressible genes were overall more consistently regulated across different experiments. This was exemplified by canonical estrogen induction (*GREB1*) or repression (*BCAS1*). In contrast, housekeeping genes such as *ACTB* exhibited a random distribution (Supplementary Fig. S2D). We identified 65 up- and 22 down-regulated genes that were enriched in top 10% percentile of altered genes and consistent across at least 50% of comparisons (Supplementary Fig. S2E). Most of these genes have previously been documented as estrogen regulated, however a few genes, such as *HEY2* and *RAB27B*, have not previously been reported as estrogen regulated (Supplementary Fig. S2E). Of note, both up- and down-regulated gene sets retain their own regulatory co-factors (e.g., MED12 as unique co-activator and EP400 as a unique co-repressor) besides ESR1/FOXA1/GATA3 as the shared nexus (Supplementary Fig. S2F). Furthermore, active histone modification marks such as H3K27ac and H3K4me3 were more enriched in E2-repressed genes loci (Supplementary Fig. S2G).

### Refining early and late estrogen response transcriptomic signatures

Transcriptomic signatures of ER action have been key to understanding endocrine therapy and resistance in breast cancer. MSigDB contains two widely-cited estrogen response signatures in the Hallmark collection, representing an early and late response(36,37). Notably, 1) both signatures were derived from only four microarray experiments and this also included an ER negative cell line MDA-MB-231 with ectopic *ESR1* overexpression; 2) only estrogen-induced but not repressed genes were included; 3) 49.5% of the genes overlap between the two signatures. Therefore, we set out to utilize the EstroGene database to derive a more representative estrogen response signature.

Among the 146 merged transcriptomic data sets, 27 different time points were annotated spanning from 5 minutes to 600 hours of estrogen stimulation. We separated all the comparisons into three signatures of duration: EstroGene_Early (< 6 hours, n=58), EstroGene_Mid (6-24 hours, n=44) and EstroGene_Late (> 24 hours, n=44) (Fig. 5A). Up- and down-regulated genes present in the top 10th percentile of regulated genes in each individual study, and consistently present across at least 50% of studies at each time period, were extracted from each signature (early, mid, and late) and intersected accordingly (Supplementary Fig. S3A). We identified 165, 59 and 136 genes representing early, mid, and late estrogen response signatures respectively (Fig. 5B and Supplementary Table S6). Intriguingly, nearly half of early response genes showed sustained estrogen regulation in the mid and late response signatures, and most mid response genes were sustained in the late response signature, hinting that more stable gene regulation might dominate after 24 hours of E2 exposure. Moreover, an early treatment duration triggered more activated than repressed genes and vice versa for late treatment duration, implicating that E2-inducible and repressible genes may entail distinct temporal regulation. We compared our EstroGene-derived estrogen response signatures with those from MSigDB. The early signatures exhibited 50% overlap whereas only 23.3% genes were found in the late signature (Supplementary Fig. S3B). Surprisingly, 3 genes (*ID2, ELF3* and *AQP3*) that are induced by E2 in the MSigDB signatures were identified as E2-repressed genes in our analysis (Supplementary Fig. S3C). Markedly, the Hallmark late signature but not the EstroGene mid or late signatures were prognostic for patients with ER+ breast cancer in METABRIC cohort, while both early response signatures remain prognostic (Supplementary Fig. S3D).

**Figure 5.**
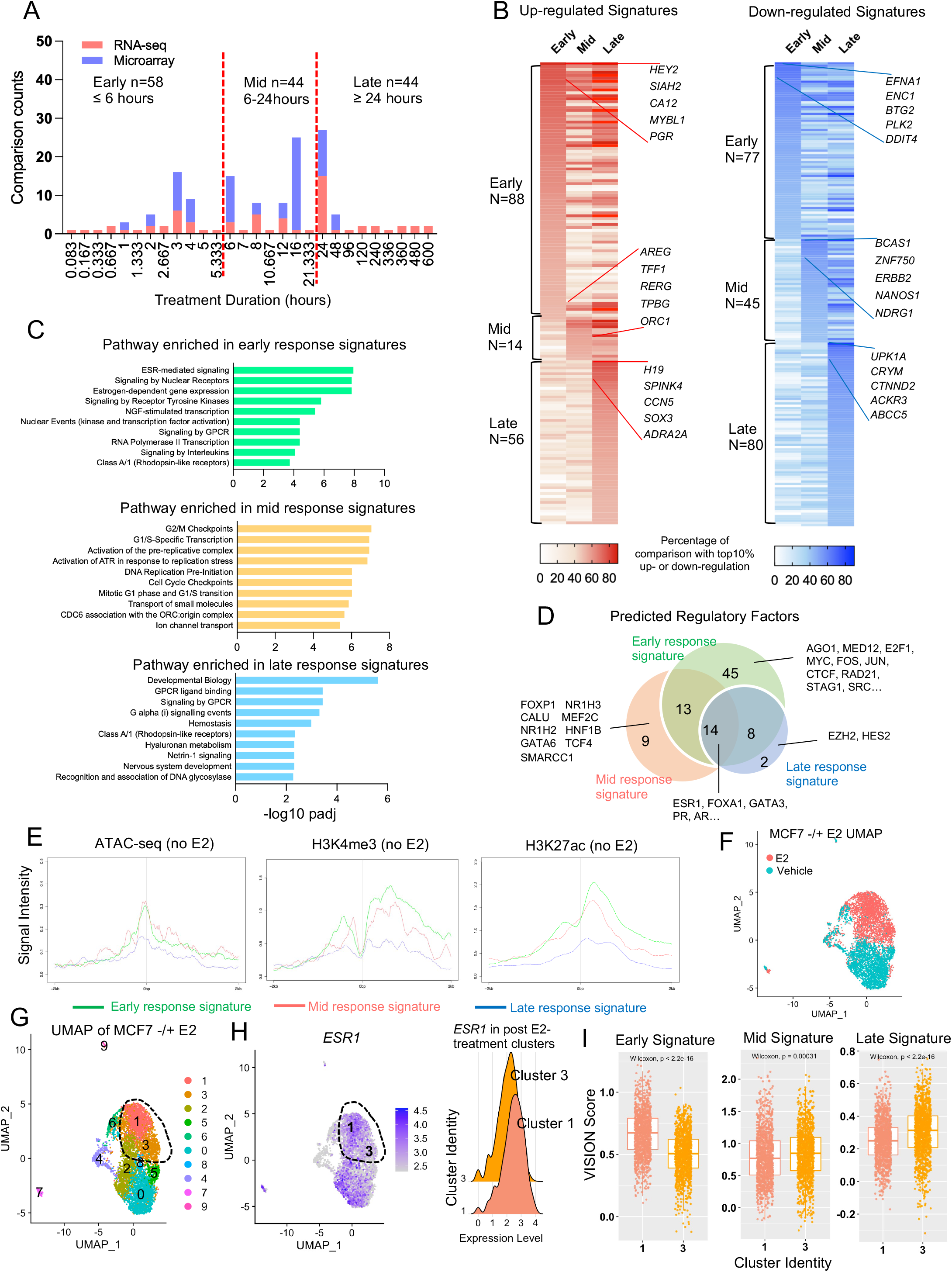
Refining early and late estrogen response transcriptomic signatures. A. Stacked histogram representing the time points of E2 treatment used in 146 transcriptomic comparisons. Three temporal courses were identified accordingly using 6 hours and 24 hours as the cutoff. B. Heatmaps illustrating the percentage falling into the top 10% percentile of up (Left panel) and down (Right panel) categories of all the genes from early, mid, and late E2 response signatures. Top five consistent genes of each category are labelled with gene names. C. Bar chart showing the significantly enriched REACTOME pathways in early, mid, and late response genes. D. Venn diagram showing the overlap of transcriptional factors predicted by LISA associated with early, mid, and late response genes. E. Intensity plot showing the signals from ChIP-seq of H3K4me3, H3K27c and ATAC-seq (no E2) on −/+ 2kb region of TSS of all early, mid, and late response genes. Epigenetic profiling data sets were downloaded from GSE99542, GSE78913 and GSE57436. F and G. UMAP showing cluster assignment under resolution 0.2 for the MCF7 single cell RNA-seq data separated by treatment (F), cluster identity (F). The post-treatment cells are circled out. H. Left panel: UMAP illustrating the expression of ESR1 in the UMAP from G. Right panel: ridge plot comparing ESR1 expressional levels in cluster 1 and 3 defined in G. I. Box plots comparing enrichment scores of early, mid, and late response signatures in each cell from cluster 1 and 3 defined in F. Mann Whitney U test was used.

Pathway enrichment analysis showed canonical ER signaling and RNA polymerase II-related transcriptional activation functions in the EstroGene_Early signature, whereas cell cycle progression was enriched in EstroGene_Mid response genes (Fig. 5C). Developmental and metabolic pathways, as well as GPCR signaling were highly enriched in EstroGene_Late response signatures (Fig. 5C). Regulatory factor prediction by Lisa(38) confirmed the ER/FOXA1/GATA3 as the central axis of gene regulation regardless of treatment duration (Fig. 5D and Supplementary Table S7). Notably, on average 5 full and 3.5 half estrogen response element (ERE) sequence were detected at the proximity (-/+ 5kb of TSS) of these E2 response genes, and they were not differentially enriched among the three classes (Supplementary Fig. S3E). Factors uniquely associated with early response genes were mainly ER cofactors as well as components of topological associated domain (e.g. CTCF, RAD21 and STAG1), consistent with previously reported E2 action on chromatin loop reprogramming(14).

Epigenetic factors such as EZH2 and SMARCC1, on the other hand, were largely enriched in mid and late response genes (Fig. 5D), in parallel with a more stable gene regulation program discerned in Fig. 5B.

We examined whether the estrogen response signatures showed a difference in chromatin accessibility of target loci in baseline conditions (i.e., no E2 stimulation). Epigenetic marks increase around −/+ 2kb of the TSS of the early and mid-estrogen response genes with more prevalently open chromatin and H3K27ac and H3K4me3 modifications than late response genes. This suggests that an initially active chromatin state may facilitate early gene stimulation events (Fig. 5E).

We further explored the heterogeneity of E2 response Using single cell RNA-seq profiling in E2-treated MCF7 cells (18). E2 stimulation explicitly separated cells into two states (Fig. 5F). We further identified two distinct clusters in the post-treatment group (Cluster 1 and 3, circled in UMAPs) differentiated by *ESR1* expression (Fig. 5G and 5H). Applying the EstroGene signatures into the data set, we found that the EstroGene_early was strongly enriched in the *ESR1*-high subcluster whereas mid and late signatures were predominant in the *ESR1*-low subpopulation.

In summary, we defined three estrogen response signatures (early, mid, and late) from 146 transcriptomic comparisons across 19 breast cancer cell lines. This integrated analysis uncovered that the timing of estrogen response is shaped by multiple levels of regulation, including unique time-dependent regulatory factors, epigenetic accessibility, and heterogeneity of ER expression.

### Identifying cell context-dependent estrogen response programs

MCF7 and T47D cell lines have been used extensively as ER+ breast cancer models. However, extrapolation of this data to breast cancer is complicated by the known heterogeneity of breast cancer and potential biases arising from cell line specific results. Importantly, while EstroGene contains transcriptomic data from 19 different breast cancer cell lines, data from MCF7 and T47D account for ~50% and ~20%, respectively, of all experiments (Fig. 6A). To characterize and describe contextual cell-line specific responses, we identified the top 10th percentile of up- and down-regulated genes in an individual study and consistent among 50% of comparisons within MCF7 or T47D experiments. For non-MCF7/T47D experiments we lowered the threshold to 40% across studies due to the larger heterogeneity in this subset (Supplementary Fig. S4A). Intersection of the three subsets yielded 89 and 96 uniquely regulated genes in MCF7 and T47D, such as *HCK* (MCF7) and *KCTD6* (T47D) (Fig. 6B, Supplementary Fig. S5B and Supplementary Table S6). We also identified 26 genes that were not regulated in MCF7 and T47D but showed E2-induction (e.g., *MCM2*) or E2-repression (e.g., *SLC12A2*) in some other cell lines (Fig. 6B and Supplementary Fig. S4B). Of note, a few targets have been reported previously as broad estrogen response targets in breast cancer such as *SGK1*(39) and *FOS* (40).

**Figure 6.**
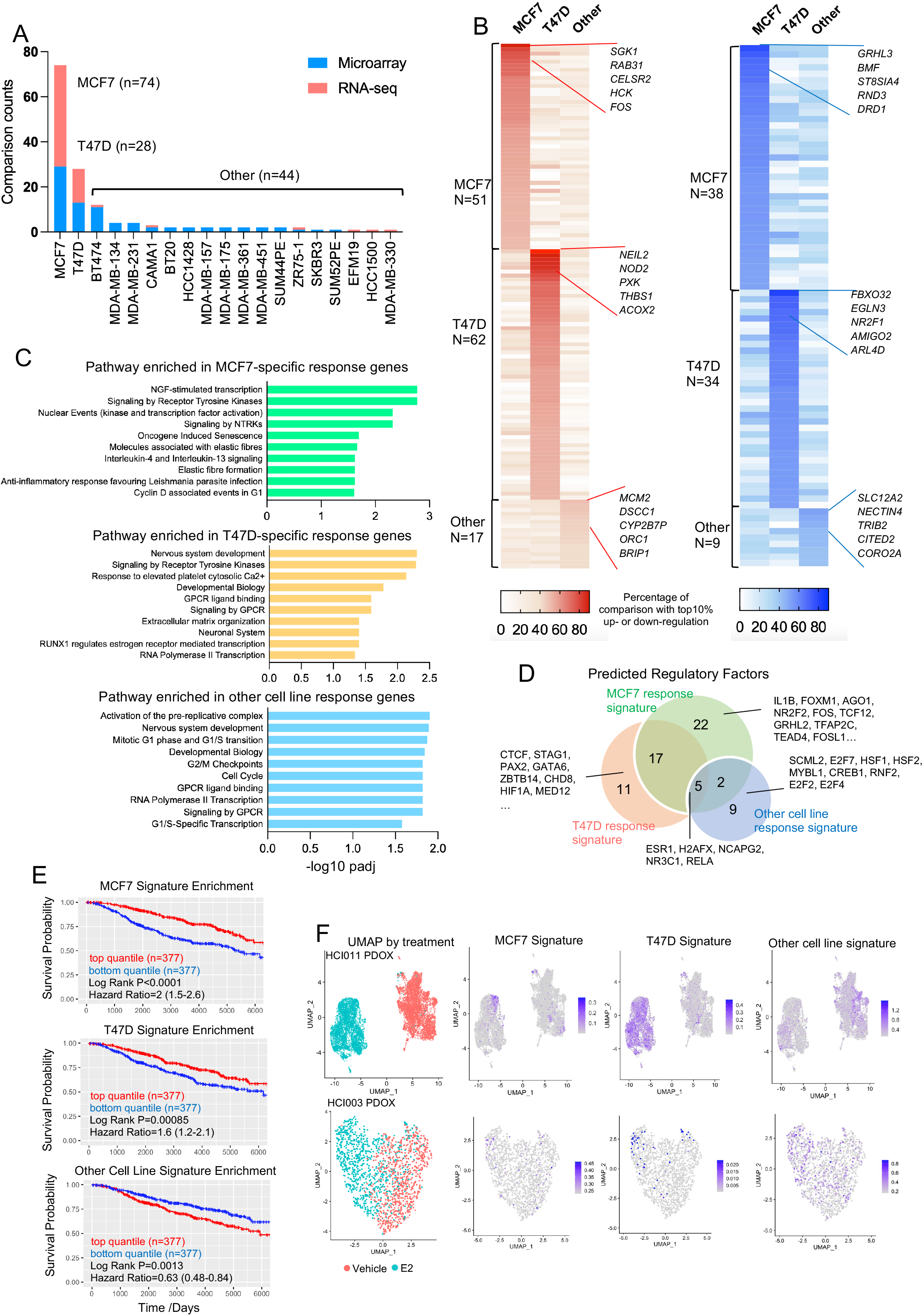
Identifying cell context-dependent estrogen response program. A. Stacked histogram representing the cell lines used in 146 transcriptomic comparisons. B. Heatmaps illustrating the percentage falling into the top 10% percentile of up (Left panel) and down (Right panel) categories of all the genes uniquely from MCF7, T47D or non-MCF7/T47D experiments. Top five consistent genes of each category are labelled with gene names. C. Bar chart showing the significantly enriched REACTOME pathways in MCF7, T47D and other cell lines uniquely E2 response genes. D. Venn diagram depicting the overlap of transcriptional factors predicted by LISA associated with MCF7, T47D and other cell lines uniquely E2 response genes. E. Kaplan-Meier plots showing the disease-specific survival (DSS) (METABRIC) comparing patients with tumors with high and low enrichment for each indicated gene sets. High and low were defined by the upper and bottom quartiles of each subset. Censored patients were labelled in cross symbols. Log rank test was used and hazard ratio with 95% CI were labelled. F. UMAP showing single cell distribution from HCI-011 (Top panel) and HCI-003 (Bottom panel) separated by treatment groups. Enrichment score of MCF7, T47D and other cell lines unique response signatures are projected on the UMAPs accordingly.

We compared pathways enrichment in cell line specific estrogen response genes. Senescence, fiber formation and inflammatory-related functions were enriched in MCF7 response genes whereas extracellular matrix, GPCR and development pathways were enriched in T47D response genes (Fig. 6C), highlighting the importance of cell line and genetic background in estrogen response. Mechanistically, MCF7 and T47D unique response genes were enriched for contextual regulatory factors (Fig. 6D and Supplementary Table S7) but showed equivalent levels of chromatin accessibility at each other’s open chromatin regions (Supplementary Fig. S4C), suggesting the unique gene induction program was due to context-dependent transcriptional regulomes rather than epigenetic changes.

We next addressed if these signatures show distinct clinical representation. Surprisingly, we found non-MCF7/T47D E2 response signature was associated with poor disease-specific survival in METABRIC ER+ cohort, while both MCF7 and T47D-specific signatures inversely correlated with good outcomes (Fig. 6E). We also calculated signature enrichment in single-cell RNA-seq data from two ER+ patient-derived xenograft organoids (HCI-003 and HCI-011) with estradiol treatment (18). We observed divergent E2 response programs related to each signature. In the HCI-003 model, the T47D-signature was homogenously increased whereas the MCF7-signature was only enriched in a small subpopulation. In contrast, in HCI-011, neither MCF7 nor T47D signatures were enriched, albeit there was a weak induction in the non-MCF7/T47D cell lines-derived signature (Fig. 6F). In conclusion, this analysis not only delineated MCF7 and T47D cell-line specific estrogen response programs, but also validated their heterogenous representations to a specific cell type within the same tumor.

### Discovering a bidirectional estrogen response program

Lastly, we examined plasticity of the directionality of estrogen response. Correlation of up- and down-regulation of all 12,429 genes in the merged data collection revealed a strong non-linear negative association, showing that most genes exhibited a single monodirectional regulation by estradiol (Fig. 7A). We also identified a subset of bidirectionally regulated genes (n=101) that are present in the top 10% of both up- and down-regulated targets and in at least 10% of comparisons (Fig. 7A and Supplementary Table S6), such as *CYP1A1, RIPOR3* and *DHRS3* (Supplementary Fig. S5A). Their divergent regulation was not associated with specific experimental conditions such as E2 treatment duration or cell line context (Supplementary Fig. S5B).

**Figure 7.**
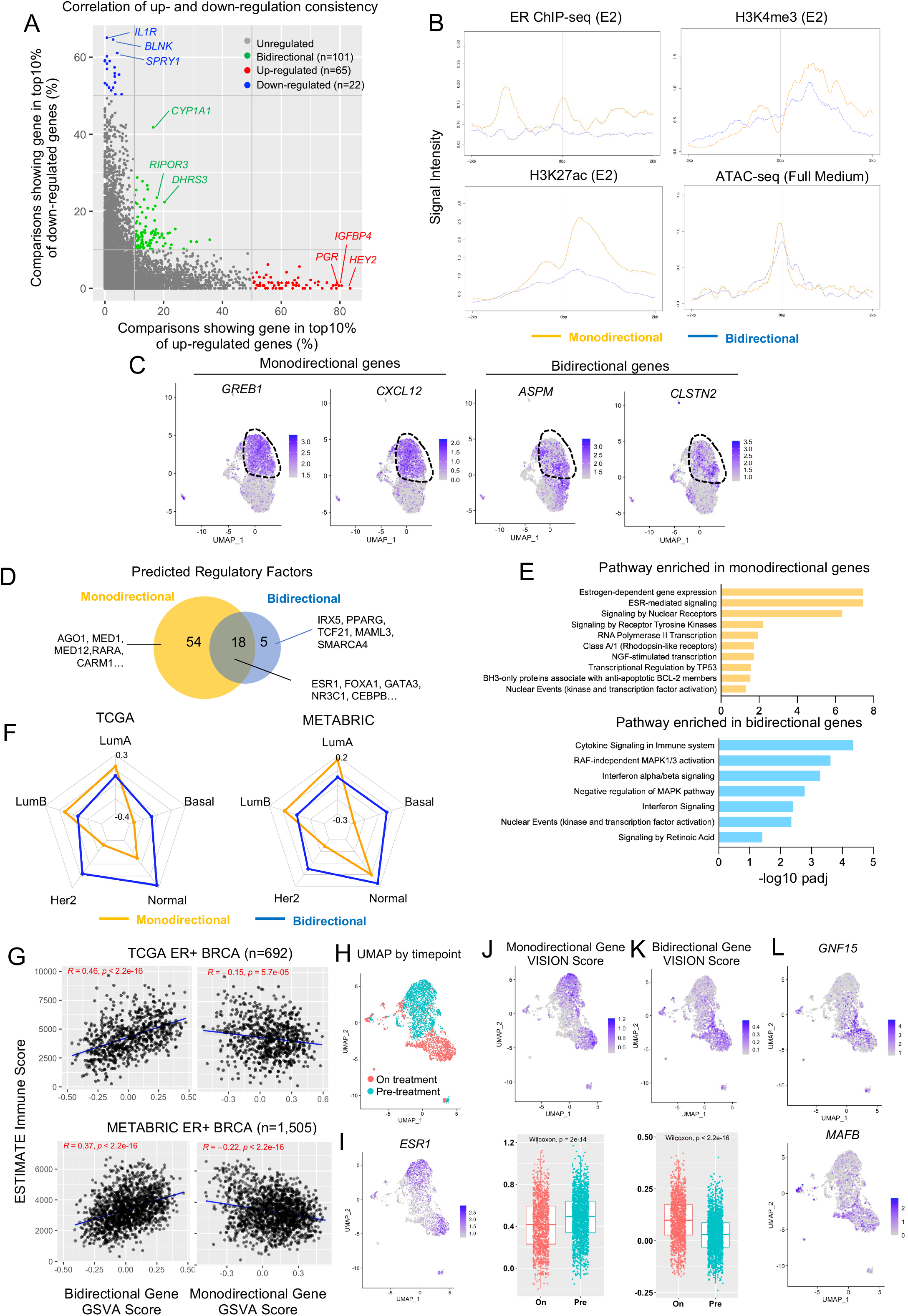
Discovering a bidirectional estrogen response program. A. Scatter plot showing the correlation of each individual gene’s percentage falling into top 10% up and down altered targets by fold changes among all 146 transcriptomic comparisons. Monodirectional genes are labelled in red (up) and blue (down). Bidirectional genes are labelled in green. B. Intensity plot showing the signals from ChIP-seq of ER (with E2), H3K4me3, H3K27ac and ATAC-seq on −/+ 2kb region of TSS of all monodirectional or bidirectional genes. Epigenetic profiling data sets were downloaded from GSE78284, GSE78913, GSE57436 and GSE102441. C. UMAP depicting the expressional levels of two representative monodirectional (*GREB1* and *CXCL12*) and bidirectional genes (*ASPM* and *CLSTN2*) are plotted on the UMAP. Post-E2-treatment cells are circled out for illustration. D. Venn diagram showing the overlap of regulatory factors predicted by LISA between monodirectional and bidirectional gene sets. E. Bar chart showing the significantly enriched REACTOME pathways in monodirectional and bidirectional gene sets. F. Radar plot showing the median enrichment scores of monodirectional and bidirectional gene signatures across PAM50 subtypes in TCGA (Left panel) and METABRIC (Right panel). G. Scatter plot depicting the Pearson correlation between enrichment scores of monodirectional (Right Panel) or bidirectional (Left panel) gene signatures with precited immune infiltration scores by ESTIMATE in ER+ tumors from TCGA (Top panel) and METABRIC (Bottom panel) cohorts. H and I. UMAP showing cancer cell distributions (H) and ESR1 gene expression (I) from two biopsies from an ER+ patient separated by anti-PD1 treatment status. J and K. Top panel: UMAP showing enrichment scores for monodirectional (J), and bidirectional (K) response genes projected on the tumor sample in H. Bottom panel: Box plot comparing enrichment scores between pre-treatment and on-treatment cancer cells. Mann-Whitney U test was applied for each comparison. L. UMAP projection of GNF15 and MAFB expression in the tumor sample described in H.

Examining cistromic and epigenetic profiles from MCF7 cells, bidirectional response genes harbored weaker ER binding and lower levels of active histone marks such as H3K4me3 and H3K27ac at their proximity (-/+ 2kb of TSS) compared to strong monodirectional response genes (Fig. 7B). Nevertheless, surrounding chromatin accessibility was not different between the two groups of genes (Fig. 7B). Using single cell RNA-seq profiling in E2-treated MCF7 cells analyzed in previous Fig. 5F (18), we found homogenous and strong enrichment of monodirectional gene signature in E2-treated cells compared vehicle-treated cells, but a bidirectional gene signature was not different between treatments (Supplementary Fig. S5C), likely due to a large number of genes which were not uniformly regulated. When examining specific genes, we observed that some bidirectional genes (e.g., *ASPM* and *CLSTN2*) were induced in a subset of cells while monodirectional genes such as *GREB1* and *CXCL12* consistently showing induction across all cells (Fig. 7C). Taken together, this analysis demonstrates that these bidirectional genes are generally transcriptionally inert and show a heterogeneous response in subpopulations of cells upon E2 exposure, indicating other cell specific factors that may be required for their regulation.

To test whether the bidirectional response is caused by specific ER regulatory co-factors possessing bivalent regulatory potential we utilized Lisa(38) and predicted 72 and 23 significantly enriched factors associated with mono- and bidirectional genes respectively (Fig. 7D). As expected, canonical factors involved in ER action such as ER, FOXA1 and GATA3 were enriched in both genes sets. Only five factors were uniquely enriched in bidirectional genes and involved pathways of NF-kB (IRX5), NOTCH (MAML3), MAPK (TCF21) signaling and SWI/SNF (SMARCA4) chromatin remodeler (Fig. 7D). Pathway enrichment analysis highlighted immune-related (e.g., cytokine signaling and interferon signaling) and MAPK pathway-relevant functions associated with these bidirectional genes (Fig. 7E). In contrast, estrogen-dependent signaling and RNA polymerase II functionals were characterized as features of monodirectional genes (Fig. 7E).

Estrogen response is a prominent feature of the ER+ luminal subtype of breast cancer. Using TCGA(41) and METABRIC(42) we compared the enrichment of mono- and bidirectional genes across PAM50 subtypes. While monodirectional genes were specifically enriched in LumA and LumB breast tumors, the bidirectional genes were not enriched in any specific PAM50 subtype but showed a slight enrichment in the normal subtype (Fig. 7F). In addition, monodirectional genes, but not bidirectional genes, were associated with prognosis in ER+ breast cancer (Supplementary Fig. S5D). Given that the pathway and TF prediction pointed to immune-related functions of these bidirectional targets, we their role in modulating immune response rather than the classic hormone-related phenotypes in breast cancer. Accordingly, we identified a strong positive correlation of predicted immune infiltration score exclusively with bidirectional genes among ER+ breast cancers in both TCGA and METABRIC cohort.

We reasoned that a subset of breast cancer cells may use the bidirectional transcriptional program as a unique stress response strategy to escape the immune surveillance. To test this hypothesis, we examined the BIOKEY cohort of single cell RNA-seq profiling in 15 intra-patient paired treatment naïve or anti-PD1 treatment ER+ breast cancer biopsies (43). Bidirectional genes were enriched in anti-PD1 treated cancers compared to pre-treatment pairs in 4/15 patients. Taking a representative patient (P17) as an example, UMAP illustrated a clear separation of all the pre- and on-treatment cells (Supplementary Fig. S5E) and we further extracted and re-clustered cancer cells by corresponding epithelial markers (Supplementary Fig. S5F, G and Fig. 7H). A monodirectional gene signature was enriched in both time points and closely linked to *ESR1* expression (Fig. 7J and 7I), whereas bidirectional genes were highly selected in on-treatment samples regardless of ER levels (Fig. 7K). This can be exemplified by genes such as *GNF15* and *MAFB* which were previously characterized for their immune suppressive roles in cancers(44,45) (Fig. 7L).

In summary, our meta-analysis identified a subset of bidirectional genes that retain both E2-inducible and repressible plasticity in breast cancer. Compared to top monodirectional regulatory counterparts, these genes show lower transcriptional inducibility, higher levels of response heterogeneity and may require unique factors for their activation. Bidirectional genes are not associated with luminal identity but rather tightly linked to immune escape particularly under immune therapy in a subpopulation of patients (Supplementary Fig. S5H).

## Discussion

The rapid growth of multi-omic cancer data poses an unprecedentedly rich resource but comes with various challenges including integration into a unified and comprehensive platform. Although several databases have preprocessed and incorporated publicly available data sets and constructed web browsers (22,23), such as Transcriptomine which focuses on nuclear receptor biology with associated metadata (24,25), no previous databases have focused on the estrogen receptor in breast cancer and merged more than one type of genome-wide platform for a high-dimensional overview. Here, we present EstroGene, a public knowledgebase, providing standardized and integrated transcriptomic and cistromic data analysis to characterize ER activation in breast cancer cells. EstroGene features curation of many E2 stimulation experiments across an extensive panel of breast cancer cell lines, E2 dose and durations with detailed experimental information abstracted from original publications. A dedicated web browser enables researchers to quickly evaluate E2 regulation of on an individual gene under defined experimental conditions and statistical cut-offs with both expression and ER proximity binding information indicating cross-data set consistency. Overall, the extensive number of datasets and methodological details we have collected allow an unprecedented opportunity to dissect the technical and biological variation in ER action.

In this study, we provide a highly practical tool, but also performed rigorous inter-data set comparisons to highlight reproducibility between studies. Both RNA-seq and ChIP-seq cross-data set analysis revealed large differences between independent data sets, and notably the overlap only included genes with strong and robust E2-induction. Pre-existing biological variation between cell lines likely play a major role in the inconsistency and lack of reproducibility between data sets. For example, a previous study using FISH identified significantly different genomic abnormalities in MCF7 cells lines from three independent institutions (46). Technical differences may also affect response. Our cross-data set analysis suggests that a longer hormone deprivation before estrogen stimulation results in stronger response to E2. Notably, our previous study revealed that components of charcoal-stripped serum vary between different manufacturers or batches, which may cause differential strengths of E2 response(47).

Finally, through hand abstraction of publications associated with public datasets, we found that key experimental details were sometimes missing such as cell line source and passage number. In addition, the method for hormone deprivation varies between studies in terms of duration and serum types (bovine vs. calf serum), which may induce additional technical variations that reduce reproducibility. This is largely in line with the challenges confronted by The Reproducibility Project, where 70% of experiments required asking authors for key reagents from the original sources(48,49). Thus, a standardized framework for experimental documentation and a reference from a centralized cell line data base such as Cellosaurus(50) is required to improve rigor and reproducibility. For instance, the cell passage number documentation may need a more uniformed recording manner in order to make it comparable across different laboratories. The EstroGene database provides the most comprehensive insight into reproducibility of studies examining ER action in breast cancer cell lines.

Estrogen response gene signatures have proven invaluable in the study of ER action in breast cancer transcriptomic datasets. Previously established early and late estrogen response signatures from MSigDB have been extensively cited in greater than 5,000 studies(36). However, studies applying these signatures rarely differentiated the biological indications between early and late ER response, partially owning to the lack of temporal specificity. Here, we derived more representative estrogen response signatures using the EstroGene database, which originated from 146 transcriptomic profiling comparisons (vs. 4 from MSigDB), 19 breast cancer cell lines (vs. 2 from MSigDB), 27 different time points (vs. 3 from MSigDB) and consisted of both activated and repressed genes (vs. activated genes only from MsigDB). Our prognostic analysis clearly reveals that only early, but not mid or late response signatures are prognostic for ER+ breast cancer patients, which yields different conclusion from the Hallmark signatures. It is plausible that endocrine therapy prominently blocks early response programs which is sufficient to suppress hormone-mediated cell growth. We hereby encourage future studies to include both Hallmark and EstroGene signatures for the analysis for a more robust and comprehensive interpretation. We identified that different rates of E2 response relate to chromatin accessible states, temporal specific TFs, and heterogeneity of ER expression. For example, the prediction of EZH2 as a unique late response gene regulator suggests that some of these genes may be indirectly induced via alteration of H3K27 methylation, or recruitment of REA at the corresponding genomic region, rather than direct ER-mediated transactivation, consistent of several earlier studies(51,52). In all, the EstroGene response signatures represent a more diverse array of ER response.

ER can trigger both transcriptional activation and repression by recruiting different cofactors(53,54). However, the plasticity of regulation upon individual genes has not been extensively explored. By merging and mining 146 E2-stimulated transcriptomic differences in multiple contexts, we unexpectedly identified a subset of genes that present as both the top E2-activated and repressed genes in different experiments. Notably, the estrogenic effects on some of these “bidirectional” targets were reported as unidirectional, as they came from a single study whereas we re-define this using meta-analysis. An example is the cytochrome P450-encoding gene *CYP1A1*, which was reported as an estrogen-repressed gene via enhanced DNA methylation following recruitment of DNMT3 in multiple breast cancer cell lines(55). The EstroGene databases shows that CYP1A1 is E2-induced in a subcollection of experiments. The mechanism behind this bivalent regulation is largely understudied and warrants future investigation. It is plausible that context-dependent and dual-function transcription factors cooperate with ER to induce divergent effects depending upon cell state and external cues. IRX5, a predicted TFs enriched in these bidirectional genes, controls downstream NF-kB signaling (56). This could either escalate or alleviate ER signaling via distinct mechanisms in different cell populations or strains depending upon culture medium component and expression levels or ER or its cofactors. Another TF factor SMARCA4, the core ATPase of the SWI/SNF complex, is enriched in bidirectional genes and may attenuate gene expression by decreasing chromatin accessibility(57) while ER might simultaneously potentiates transcription of these genes. By mining single-cell RNA-seq profiling of series biopsies from an anti-PD-1 treated breast cancer cohort, we further found that the E2 response plasticity might be used by cancer cells to facilitate their escape from immune surveillance, while it may not affect endocrine therapy outcomes. For example, the induction of *GDF15* with anti-PD-1 exposure could largely cause immune suppression via CD44-mediated suppression of dendritic cells maturation(58) and blockade of cytotoxic T cell recruitment(59). This is consistent with a previous report describing the role of ER signaling in suppressing cancer immune response(60). Due to the limited sample analyzed here, the clinical association will need to be strengthened in a larger cohort in the future.

The EstroGene database shows that MCF7 and T47D cells account for 70% of publicly available E2-regualted data sets. This raises a concern about the bias of models and generalized interpretability of findings. Consistent with this, we interrogated cell specific effects which may be mediated by context-dependent transcriptional factors. For example, unique upregulation of FOS in MCF7 could trigger a secondary transcriptional cascade via Jun/Fos signaling. In parallel, CHD8, a required epigenetic factor to activate progesterone receptor-dependent enhancers(61), is exclusively enriched in PR positive T47D cell lines. Consistent with this, our previous work introducing constitutively activated estrogen receptor mutations into MCF7 and T47D cell lines revealed divergent transcriptomic reprogramming and context-dependent metastatic phenotypes (62,63). Our results also suggested that the association of E2 response signature enrichment degree and patient survival outcome with endocrine therapy are context-dependent. Some E2 response genes within non-MCF7/T47D cell line may also propagate other essential steps of tumor progression such as immune escape and metastatic spread and hence correlates to poorer survival outcome. The contextual E2 response gene modules produced here offers a useful resource helping researchers to potentially avoid selection of biased targets for in-depth characterizations. The growing utility of new generation breast cancer models such as patient-derived organoids is indispensable to preserve the heterogenous nature of breast cancer in the future(64). However, in vitro culture can still introduce undesired variabilities that impact the physiologic relevance of the findings and thus in vivo validation is of utmost importance.

In conclusion, the EstroGene database is a user-friendly platform for analysis and visualization of ER regulated gene expression. We intend to extend this platform to include further data sets such as ATAC-seq and Hi-C for more extensive mechanistic insight. We also plan to incorporate data sets from breast cancer models harboring clinically relevant estrogen receptor variants such as hotspot mutations and fusions and with anti-ER agent’s treatments to yield consensus of ER regulomes associated with endocrine resistance. We also expect to continue to ingest and process further datasets into the EstroGene browser with continuous crowdsourcing from the research community. We hope the EstroGene database will ultimately support global cancer research and beyond.

## Materials and Methods

### Data curation and documentation

To obtain a harmonized estrogen receptor related base in breast cancer, we established a standardized curation model with three main steps. First, we conducted a literature search from the Gene Expression Omnibus (GEO database) using the combination of “estrogen” or “E2” or “estradiol” plus “breast cancer” plus the name a specific type of sequencing technology (e.g., “RNA-seq” or “RNA-sequencing”) towards publications released earlier than January 2022. Secondly, we manually reviewed these articles, only literatures conducting E2 stimulation experiments on human breast cancer cell lines were incorporated into EstroGene database. We curated details of publications and experimental designs including cell models, E2 dose, duration, control type, culturing medium, hormone deprivation methods, estradiol product and dissolvent information, library preparation method and NGS sequencing platforms. All the relevant information is summarized in Supplementary Table S1. An additional proof reading was performed by an independent researcher from our team to ensure the accuracy of our documentation. In addition, we posted our platform and the metadata table of all the curations online via Twitter in October 2022 for continuous crowdsourcing with proper instructions for new data set notification. Data curation does not involve any bias reduction techniques.

### Webserver construction and implementation

The EstroGene database Application uses MySQL (https://www.mysql.com/) and Django (https://www.djangoproject.com/) Framework to manage request from frontend webpage. The front end utilizes Javascript to dynamically render the webpage. In particular, jQuery https://jquery.com/) and Ajax (https://developer.mozilla.org/en-US/docs/Web/Guide/AJAX) are deployed to support the core features of the EstroGene database. jQuery is a fast, small, and feature-rich JavaScript library that makes it easy to manipulate the Document Object Model (DOM), handle events, and perform HTTP requests from Ajax calls. It is designed to simplify the process of writing JavaScript code and makes it easier to work with web pages. Additionally, CharJS (https://www.chartjs.org/) is also used to enable interactive charts and visualization.

### Transcriptomic data process and analysis

For RNA-seq data sets, we uniformly downloaded 375 raw fastq files from the corresponding data sets from GEO with the SRR accession numbers. We used Salmon v0.14.1(65) to align the reads to hg38 reference genome (Genecode.v29) and genes counts export using Tximport assignment on EnsDB.Hsapienes.v86. Genes with constantly 0 counts were removed and DESeq2(66) was used to compute Log2Fold Change and adjust p values of each gene between control and E2 stimulated samples. For specific data sets lacking replicates, we generated Log2 fold change of each gene by subtracting TMM normalized Log2(CPM+1) values of controls from the corresponding stimulated samples.

For microarray data sets, we collected the raw array files from GEO database and normalized the data with different packages according to the platform. Affy(67) and oligo(68) packages were used to process Affymetrix-based microarray data following RMA normalizations. For illumina-based microarray data, lumi(69) package was used for data normalization. For data generated based on Agilent platform, loess normalization was performed directly on preprocessed data were downloaded from GEO. Different version of probe ID were converted to gene ID using BioMart(70) package. Probes representing the same gene were merged by averaging the normalized intensity. Limma(71) was used to compute differential expressing genes for data sets including biological replicates. For experiments without replicates, log2 fold changes were calculated by subtracting the control values from the matched E2 treated samples.

For clinical sample analysis, TCGA RNAseq reads were reprocessed using Salmon v0.14.1(65) and Log2 (TPM+1) values were used. For the METABRIC data set, normalized probe intensity values were obtained from Synapse under license to AVL. For genes with multiple probes, probes with the highest inter-quartile range (IQR) were selected to represent the gene. Batch effects of the four RNA-seq experiments(28–31) (GSE73663, GSE51403, GSE56066 and GSE78167) were removed using “removeBatchEffect” function of “limma(71)” package. Gene set variation analysis was performed using “GSVA” package(72). Survival comparisons were processed using “survival” and “survminer” packages(73) using Cox Proportional-Hazards model and log-rank test. Data visualizations were performed using “ggpubr(74)” “fmsb(75)” and “VennDiagram(76)”. Gene set enrichment analysis was performed using the “investigation” function from the MSigDB webserver using the REACTOME gene set collection with FDR below 0.05.

For single-cell RNA-seq data analysis, raw read counts matrix and metadata were downloaded from http://biokey.lambrechtslab.org./ for the BIOKEY cohort(43) and GSE154873 for E2 stimulated scRNA-seq of MCF7, HCI-003 and HCI-011 models(18). Seurat objects were created using Seurat (version 4) package for further analysis(77). Genes with detected expression in less than 3 cells, as well as cells expressing less than 500 genes or containing more than 20% mitochondrial genes were removed, resulting in 6,439 (MCF7), 1,615 (HCI-003), 13,470 (HCI-011) and 3,258 cancer cells out of 6,391 total cells from patient#17 for the BiOKEY cohort. Mitochondrial genes and cell cycle scores were regressed out before principal component analysis, and a shared nearest neighbor optimization-based clustering method was used for identifying cell clusters. VISION package was used to assign enrichment scores of each signature to each single cell(78). Log normalized counts values genes or VISION score were visualized using “FeaturePlot” function.

### ChIP-seq data process and analysis

ChIP-seq raw fastq files were downloaded from GEO with corresponding SRR accession numbers. Reads were aligned to hg19 genome assembly using Bowtie 2.0 (79), and peaks were called using MACS2.0 with q value below 0.05 (80). Quality control was conducted using ChIPQC package(81). We used DiffBind package (82) to perform principal component analysis, identify gained and lost peaks by intersect BED files. For ER ChIP-seq from C22 with two biological replicates, we first derived the consensus peaks between each group’s replicates and then overlap control and E2 treatment groups. Intensity plots for binding peaks were visualized by Seqplots(83) using BigWig files and Bed files as input. Annotation of genes at peak proximity and genomic feature distribution was conducted using ChIPseeker (84), taking the promoter region as +/-3000 bp of the transcriptional start site (TSS) and 50kb as peak flank distance. For motif enrichment analysis, fasta sequences were extracted from each genomic interval using bedtools(85) and ERE motif enrichment was calculated using the AME module from the MEME Suite(86). For integration of other epigenetic data, pre-processed BigWig files for ATAC-seq from MCF7 and T47D cells (GSE99542, GSE102441 and GSE84515)(87,88), FAIRE profiling (GSE25710)(89), ChIP-seq for H3K27ac (GSE78913)(90), H3K4me3 (GSE57436)(91) and H3K9me3 (GSE96517)(92) were downloaded from the Cistrome DB. Conversion of BED file between hg19 and hg38 reference genome was conducted using lift genome annotation function from ucsc browser (https://genome.ucsc.edu/cgi-bin/hgLiftOver) before integration.

## Data Availability

Details of all the curated 136 data sets are summarized in Supplementary Table S1. This includes all the associated publication information, GEO accession numbers, experimental designs including cell models, E2 dose, duration, control type, culturing medium, hormone deprivation methods, estradiol product and dissolvent information, library preparation method and NGS sequencing platforms.

For the inter-study concordance analysis, detailed information of four RNA-seq and four ER ChIP-seq data sets are summarized in Supplementary Table S2 and S4 respectively. RNA-seq data and clinical information from TCGA and METABRIC were obtained from the GSE62944 and Synapse software platform under accession number syn1688369, respectively.

For integration of other epigenetic data, pre-processed BigWig files for ATAC-seq from MCF7 and T47D cells (GSE99542, GSE102441 and GSE84515), FAIRE profiling (GSE25710), ChIP-seq for H3K27ac (GSE78913), H3K4me3 (GSE57436) and H3K9me3 (GSE96517) were downloaded from the Cistrome DB.

For single-cell RNA-seq data analysis, raw read counts matrix and metadata were downloaded from http://biokey.lambrechtslab.org./ for the BIOKEY cohort and GSE154873 for E2 stimulated scRNA-seq of MCF7, HCI-003 and HCI-011 models.

## Acknowledgement

Yang Wu was a former visiting research scholar at the University of Pittsburgh School of Medicine supported by funds from The China Scholarship Council and Tsinghua University. The authors would like to thank Dorothy Carter for her technical assistance in this study.

