## Supplementary Figures for "EstroGene database reveals diverse temporal, context-dependent and directional estrogen receptor regulomes in breast cancer"

### Supplementary Figure S1

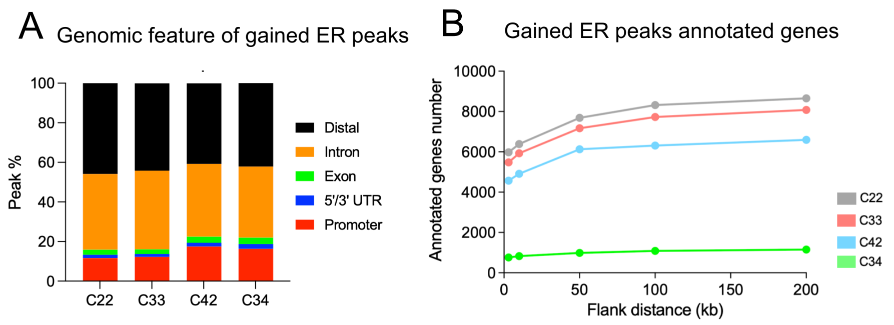


**Supplementary Fig. S1. (Related to Fig. 4)**
A. Stacked plots showing the genomic feature distributions of gained ER peak derived from four independent experiments.
B. Line plot illustrating the annotated gene numbers with increasing flank distance for associated gene calling of the gained ER peaks from the four studies.

### Supplementary Figure 2

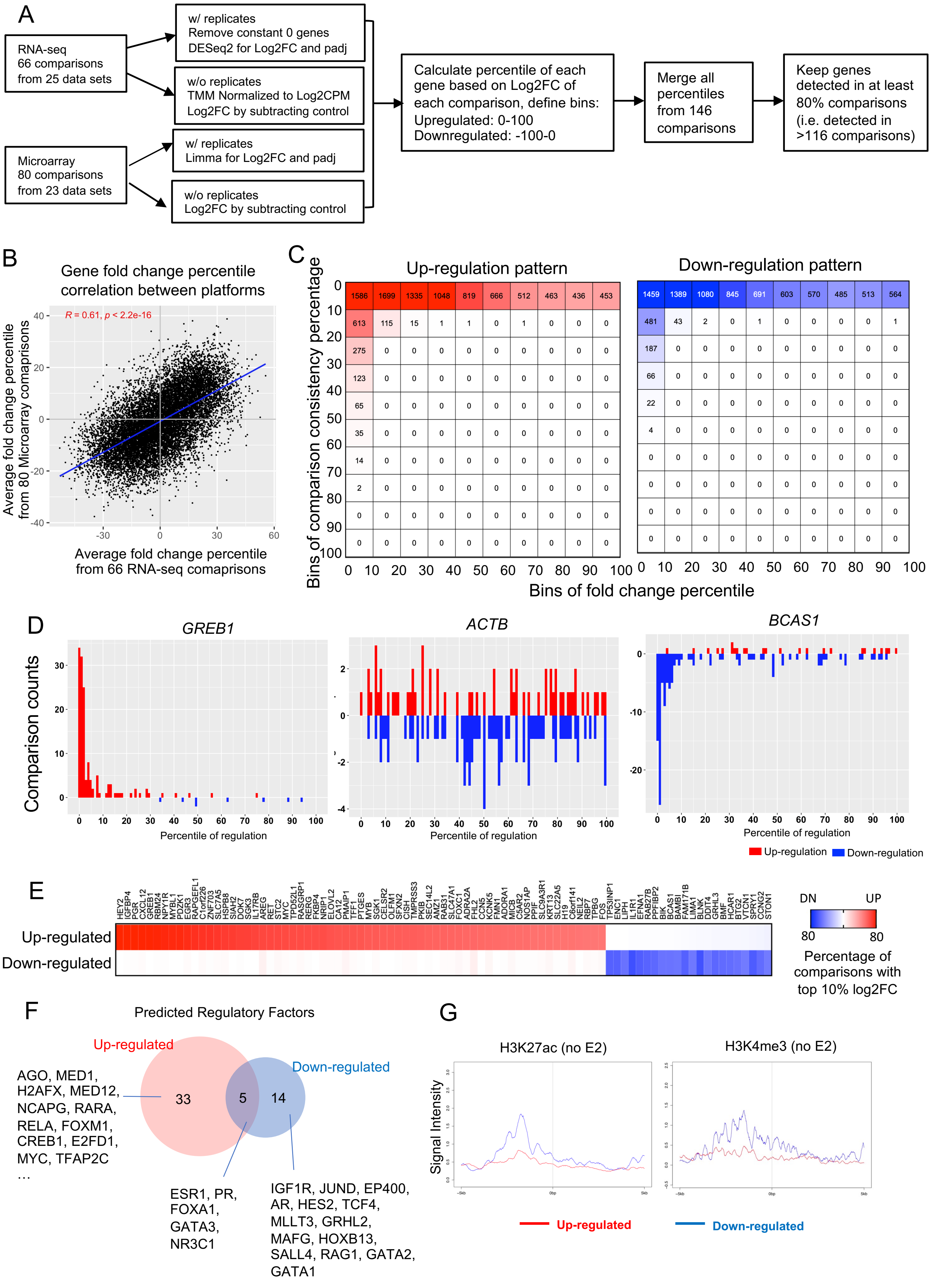

**Supplementary Fig. S2-Continued (Related to Fig. 5)**

A. A flow chart illustrating the procedure of merging 146 transcriptomic comparisons by percentile-based method.

B. Scattered plot depicting the average fold change percentile correlation of each gene between RNA-seq and microarray platforms. P value was calculated based on Pearson correlation.

C. Heatmaps showing the number of up (Left panel) and down (Right panel) regulated genes in different fold change percentiles and consistency percentages across all the comparisons.

D. Histograms showing comparison counts of each one percentage bin of regulation towards an E2-induced gene *GREB1*, E2-repressed gene *BCAS1* and non-E2 regulated gene *ACTB*.

E. Heatmap summarizing the 65 and 22 up and downregulated genes that fall in top 10% altered targets by fold changes and consistent in at least 50% of comparisons.

F. Venn diagram showing the overlap of regulatory factors predicted by LISA between highly consistent E2-up- and down-regulated gene sets shown in D.

G. Intensity plot showing the signals from ChIP-seq of H3K4me3 and H3K27ac (no E2) on -/+5kb region of TSS of all E2-up- and down-regulated genes. Epigenetic profiling data sets were downloaded from GSE78913 and GSE57436.

### Supplementary Figure 3

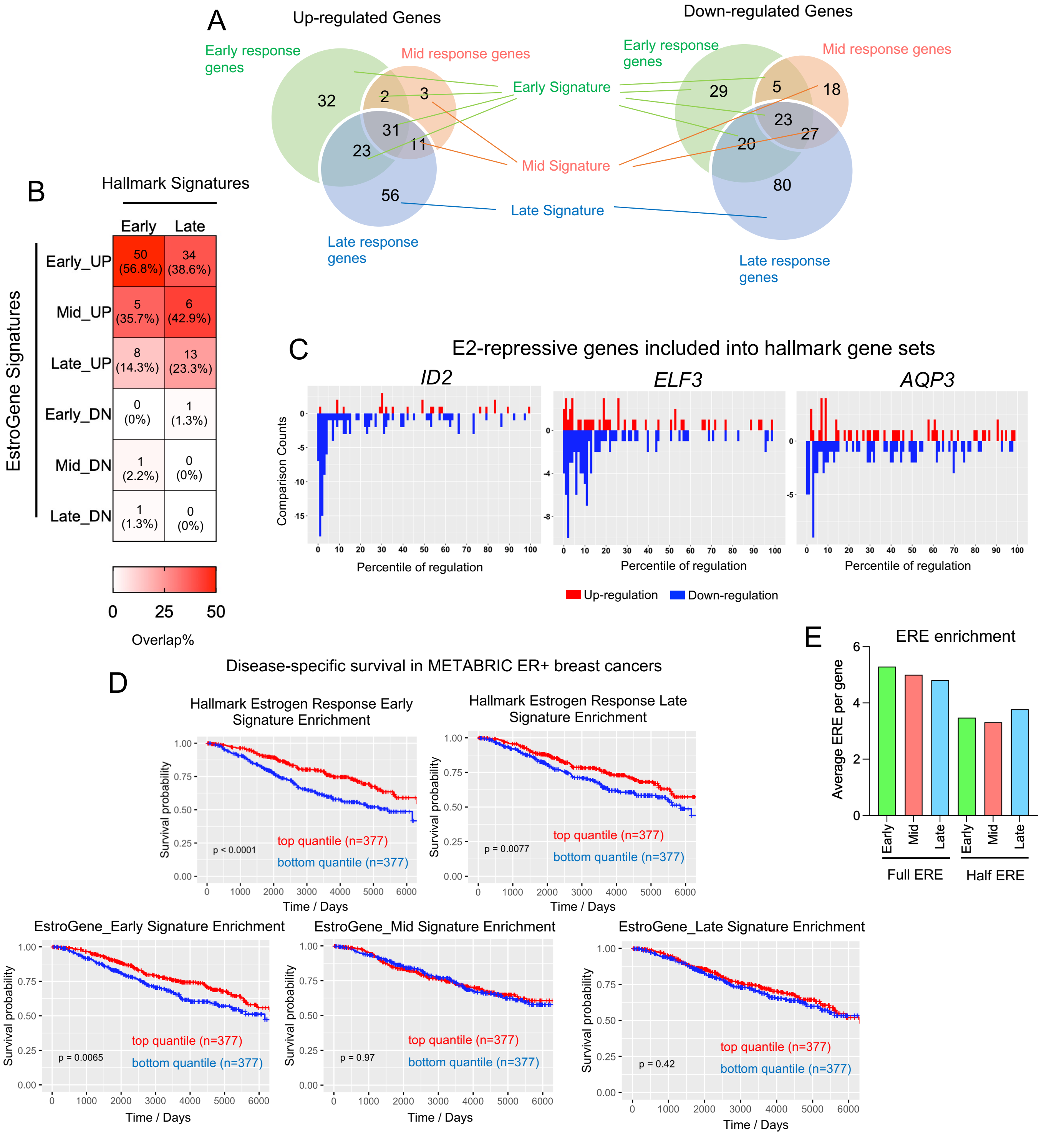


**Supplementary Fig. S3 (Related to Fig. 5)**

A. Venn diagrams demonstrating the overlap between E2 response genes extracted from comparisons of early (<6h), mid (6-24 h) and late (>24h) time points with top 10% percentile alterations and consistent across at least 50% of each comparison sub-collection.

B. Heatmap summarizing the overlap ratio of Hallmark Estrogen Response Early and Late signatures with early, mid and late signatures (separated by up and down) generated from this study.

C. Histograms showing comparison counts of each one percentage bin of regulation towards three E2-repressive genes included into the Hallmark Estrogen Response signatures.

D. Kaplan-Meier plots showing the disease-specific survival (DSS) (METABRIC) comparing patients with tumors with high and low enrichment for each indicated gene sets. High and low were defined by the upper and bottom quartiles of each subset. Censored patients were labelled in cross symbols. Log rank test was used.

E. Histogram showing the full ERE and half ERE motif enrichment per gene in EstroGene Early, Mid and Late subgroups.

### Supplementary Figure 4

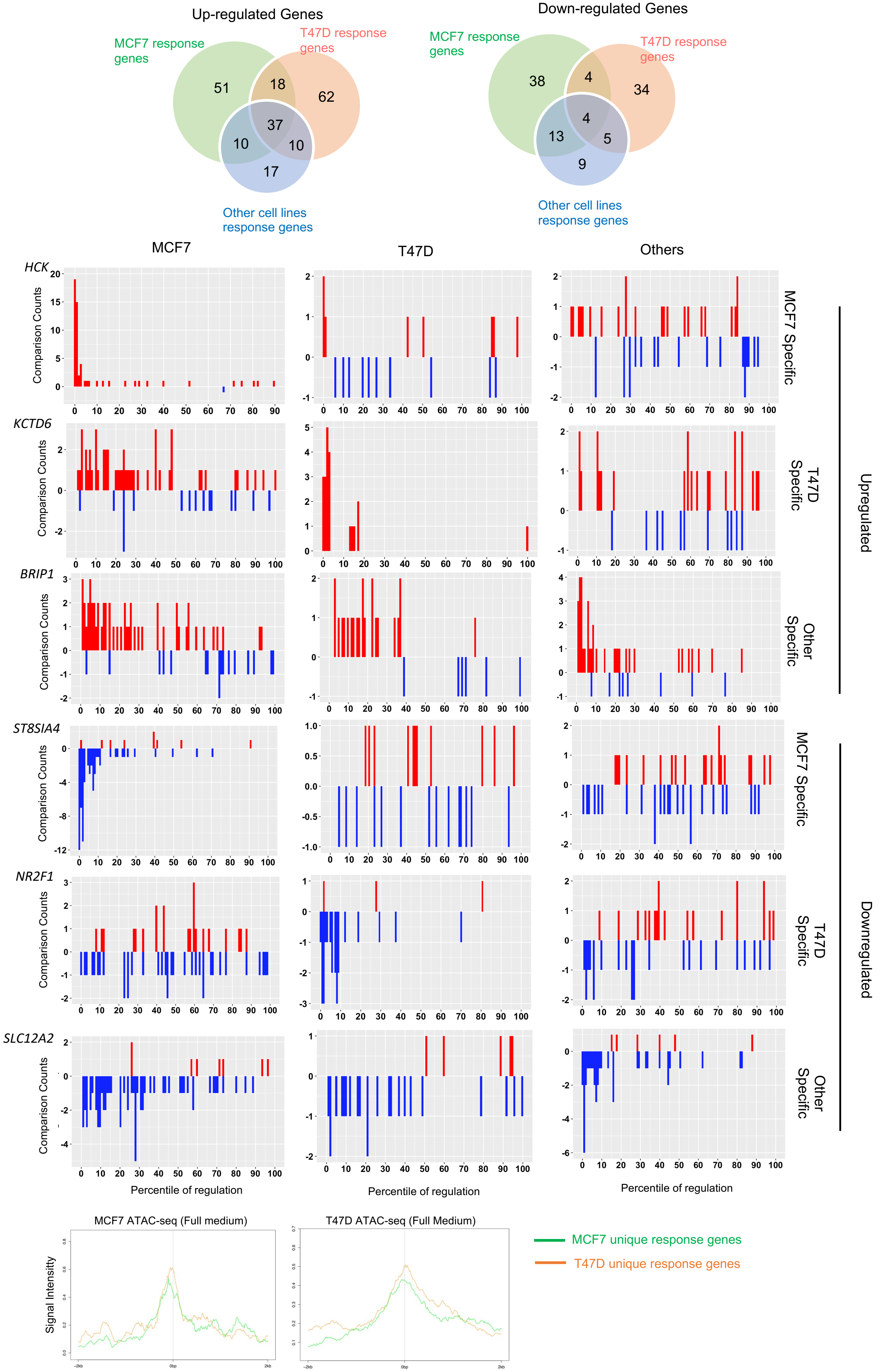

**Supplementary Fig. S4 (Related to Fig. 6)**

A. Venn diagrams demonstrating the overlap between E2 response genes extracted from comparisons limited to MCF7, T47D and non-MCF7/T47D experiments with top 10% percentile alterations and consistent across at least 50% of each comparison sub-collection.

B. Histograms showing comparison counts of each one percentage bin of regulation towards representative up- and down-regulation genes identified from each context.

C. Intensity plot showing the signals from ATAC-seq of MCF7 and T47D cells on -/+ 2kb region of TSS of MCF7 and T47D-unique E2 response genes. Epigenetic profiles are downloaded from GSE102441 and GSE84515.

### Supplementary Figure 5

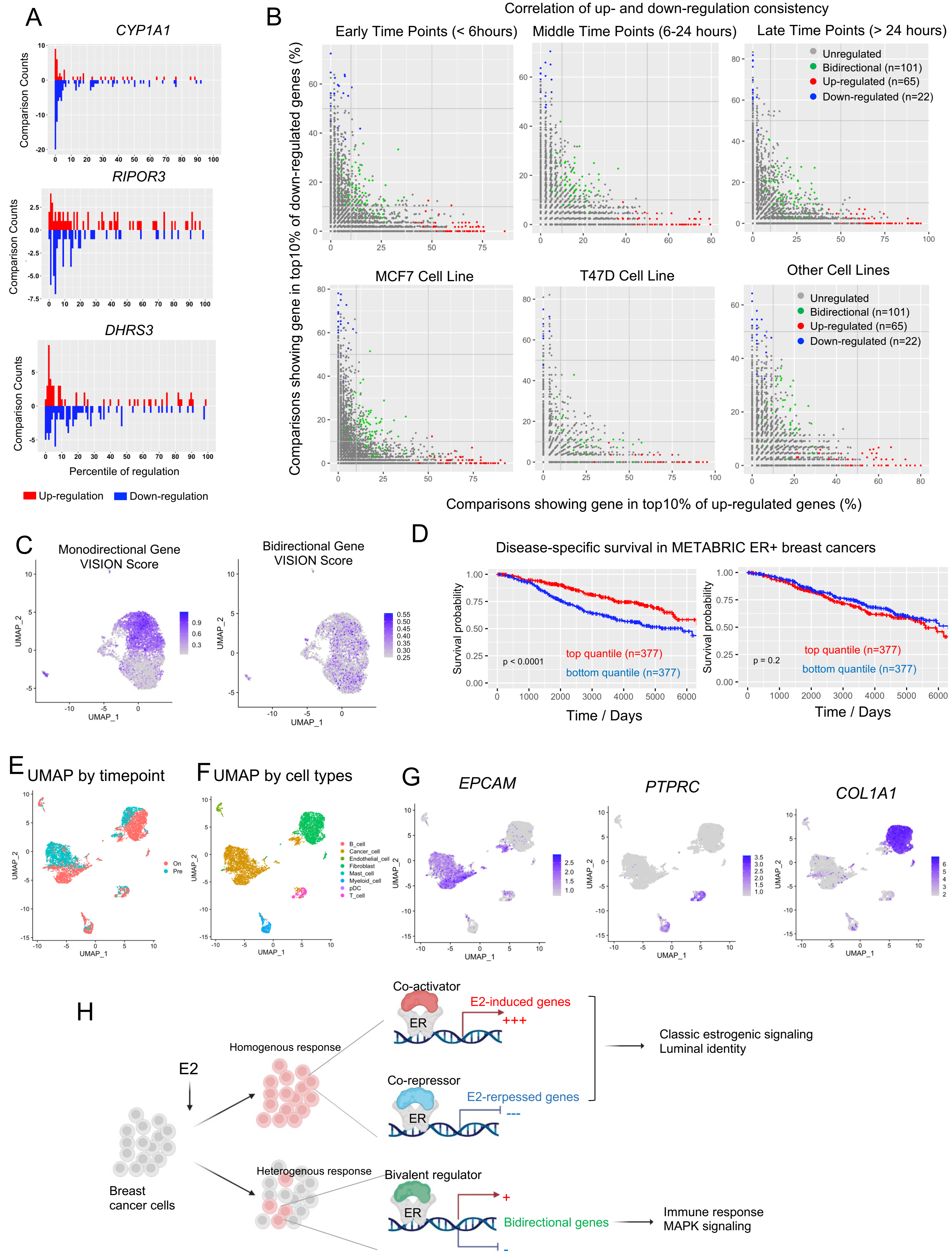

**Supplementary Fig. S5-Continued (Related to Fig. 7)**

A. Histograms showing comparison counts of each one percentage bin of regulation towards three representative bidirectional E2 response genes *CYP1A1*, *RIPOR3*, *DHRS3*.

B. Scattered plot showing the correlation of each individual gene's percentage falling into top 10% up and down altered targets by fold changes comparisons limited to specific time courses (Top panel) and cell lines (Bottom panel). Monodirectional genes are labelled in red (up) and blue (down). Bidirectional genes are labelled in green.

C. UMAP showing enrichment score of mono- and bidirectional E2 response genes in the MCF7 single cell RNA-seq data sets described in Fig. 5C.

D. Kaplan-Meier plots showing the disease-specific survival (DSS) (METABRIC) comparing patients with tumors with high and low enrichment for monodirectional or bidirectional gene sets. High and low were defined by the upper and bottom quartiles of each subset. Censored patients were labelled in cross symbols. Log rank test was used and hazard ratio with 95% CI were labelled.

E and G. UMAP showing all cell distributions (E), cell subtype assignment (F) and epithelial (EPCAM), lymphocytes (PTPRC) and fibroblast (COLA1A) marker expression (G) from two biopsies of an ER+ patient separated by anti-PD1 treatment status from the BIOKEY cohort.

H. A schematic illustration for mechanistic and functional perspectives regarding to mono- and bidirectional estrogen response genes.
